# Sassy: Fuzzy Searching DNA Sequences using SIMD

**DOI:** 10.1101/2025.07.22.666207

**Authors:** Rick Beeloo, Ragnar Groot Koerkamp

## Abstract

**Motivation:** Approximate string matching (ASM) is the problem of finding all occurrences of a pattern in a text while allowing up to *k* errors. Many modern methods use *seed-chain-extend*, which is fast in practice, but does not guarantee finding *all* matches with ≤ *k* errors. However, applications such as CRISPR off-target detection require exhaustive results.

**Methods:** We introduce Sassy, a library and tool for ASM of short patterns in long texts. Sassy splits the text into 4 parts that are searched in parallel, and uses bitvectors in the text direction rather than the pattern direction. This has compexity *O*(*k*⌈*n/W* ⌉) when searching a random text of length *n*, where *W* = 256 is the SIMD width, and provides significant speedups for small *k*.

Separately, we allow matches of the pattern to extend beyond the text for an *overhang cost* of e.g. *α* = 0.5 per character, to find matches near contig or read ends.

**Results:** Sassy is 4× to 15× faster than Edlib for patterns ≤ 1000bp, and can search text with a throughput near 2 Gbp/s. Likewise, Sassy is over 100× faster than parasail. We apply Sassy to CRISPR off-target detection by searching 61 guide sequences in a human genome. Sassy is 100× faster than SWOffinder and only slightly slower (for *k* ≤ 3) than CHOPOFF, for which building its index takes 20 minutes. Sassy also scales well to larger *k*, unlike CHOPOFF whose index took over 10 hours to build for *k* = 5.

**Availibility:** Sassy is available as library and binary at https://github.com/RagnarGrootKoerkamp/sassy, and archived at swh:1:dir:e884758dce5777a441bc2799dc8824e563c5f97b.

## 1. Introduction

Approximate string matching (ASM) is the problem of finding all matches of a pattern *P* of length *m* in a text *T* of length *n* with at most *k* errors [Navarro, 2001]. In this paper, we consider errors under the unit-cost edit distance, also known as Levenshtein distance [Levenshtein et al., 1966]. ASM has applications in many different fields. Specifically in bioinformatics, instances of ASM are CRISPR off-target detection [Yaish et al., 2024, Roux et al., 2025] and searching barcodes for demultiplexing [Cheng et al., 2024, Beeloo et al., 2025].

Recent years have seen a large number of papers on speeding up the related problem of (semi-)global alignment by using faster implementations (bitpacking, SIMD), faster algorithms (A*), and better banding heuristics (see Section 1.2). Simultaneously, there is a lot of research on *mapping*: aligning, say, 1 kbp reads against static text indices that can range from megabases to gigabases in size, without the guarantee of finding *all* matches. Since this guarantee is important for many bioinformatic applications, we identify that there is no modern, SIMD-based tool for ASM. `Sassy` (<u>S</u>IMD <u>A</u>pproximate <u>S</u>tring <u>S</u>earcher) fills this gap.

### 1.1 Contributions

`Sassy` is a conceptually simple but highly efficient command line tool and Rust library for approximate string matching. `Sassy` targets patterns with length up to around a thousand characters. It supports both ASCII and (IUPAC) DNA sequences, runs on both AVX2 (x86-64) and NEON (ARM), and comes with C and Python bindings. The underlying algorithm does not require a precomputed text index and instead can operate directly on the records of an input stream. This makes it especially suitable for e.g. searching a pattern while streaming DNA reads, one-off searches in assembled genomes, and reference-free analysis.

On a high level, our main contributions are:

1. We argue that while similar, semi-global alignment, mapping, and ASM are all distinct problems, and that for certain applications, exact methods for ASM are required and currently not available.
2. We define what it means to “report all matches”, and choose to report only local minima by default. We note that reverse-complementing inputs can give different results (Figure 2).
3. We develop an efficient implementation of ASM. Algorithmically this has two small novelties: 1) bit-packing in the text direction, rather than the pattern direction, and 2) intra-sequence parallellism by splitting the text into 4 chunks that are processed in parallel using *W* = 256 bit SIMD. This leads to expected-case complexity *O*(*k*⌈*n/W* ⌉) when matching against random text, and *O*(*m*⌈*n/W* ⌉) in the worst-case when excluding the time for tracebacks.
4. In Appendix B, we introduce an *overhang cost α* = 0.5 that allows and controls the cost of *overhanging* alignments extending beyond the text.
5. `Sassy` is 4 − 15× faster than Edlib for patterns up to length 1000 with up to 5% divergence.
6. `Sassy` is 100× faster than Swoffinder for CRISPR off-target detection, and equally fast or faster than the index-based CHOPOFF while reporting identical matches.

### 1.2 Previous work

ASM has been extensively studied between 1980 and 2000, mostly concluding in the bit-packing algorithm of Myers [1999]. For word size *w* = 64, this has worst-case complexity *O*(⌈*m/w*⌉*n*), or expected-case complexity *O*(⌈*k/w*⌉*n*) on random text. Since then, research has shifted to other types of pairwise alignment. Indeed, both *global alignment* and *mapping* are very active areas of research on similar but slightly different problems. Some methods developed for those problems can also be applied to ASM. Unfortunately, they usually do not guarantee to return all matches, either because they only return best matches, as in *semi-global alignment*, or because of their heuristic nature in case of mappers. Before discussing the older results on ASM itself in detail, we first cover some recent work on these related problems, so that the differences can be appreciated.

#### Global alignment

In *global alignment*, the pattern *P* is aligned against the *entire* text *T*. The lengths *m* := |*P* | and *n* := |*T* | are typically relatively close to each other, and may range from tens to millions of bases. The classical Needleman-Wunsch [Needleman and Wunsch, 1970] (or Wagner-Fischer/Levenshtein [Wagner and Fischer, 1974, Levenshtein et al., 1966]) algorithm requires *O*(*nm*) time and space, although space can be reduced to *O*(min(*m, n*)) when only the alignment cost is needed. While no algorithm breaks the worst-case *O*(*n*^2*−ε*^) barrier under SETH [Backurs and Indyk, 2015], many practical methods achieve sub-quadratic performance on typical inputs. For instance, Ukkonen’s band-doubling method [Ukkonen, 1985b] runs in *O*(*ns*) time, where *s* is the edit distance, and diagonal-transition approaches [Myers, 1986, Ukkonen, 1985a] attain *O*(*n* + *s*^2^) both in expectation on random texts and in practice. A recent implementation of this (for affine costs) is in WFA [Marco-Sola et al., 2021] and its extension BiWFA [Marco-Sola et al., 2023] with reduced memory usage. Another key technique in accelerating global alignment is bit-packing, pioneered by Myers in 1999 [Myers, 1999]. Rather than processing each DP cell individually, cost differences can be stored in word-size *w* = 64 bit vectors, allowing the processing of 64 DP states at once. This reduced the time complexity to *O*(*n*⌈*m/w*⌉), or *O*(*n*⌈*s/w*⌉) with banding. This bit-packing is implemented in the commonly used tool Edlib [Šošić and šikić, 2017]. Modern CPUs can process more than 64 bits in SIMD registers (e.g. 256 bits for AVX2 or 128 bits for NEON). To effectively use parallelization, the DP matrix is often broken into smaller regions [Farrar, 2006, Wozniak, 1997, Liu and Steinegger, 2021], allowing parallel processing such as in KSW2 [Li, 2018, Suzuki and Kasahara, 2018], BSAlign [Zhang et al., 2019], and SeqMatcher [Espinosa et al., 2024]. Additionally, unlike for ASM, heuristics such as X-drop can be employed to reduce the search space [Altschul et al., 1990, Suzuki and Kasahara, 2018, Liu and Steinegger, 2021, Walia et al., 2024], while losing the guarantee that the best alignment is found. QuickEd [Doblas et al., 2025] first computes an approximate banded alignment and uses this as *R. Beeloo and R. Groot Koerkamp* input for an exact alignment. A*PA and A*PA2 instead bound the search region by using A*, and retaining the guarantee that an optimal alignment is found [Groot Koerkamp and Ivanov, 2024, Groot Koerkamp, 2024].

#### Semi-global alignment

In *semi-global* alignment, a pattern *P* is aligned to a *substring* of a longer text *T*, and gaps at the start and end of *T* do not incur a penalty. Like in global alignment, only the alignment(s) with the lowest number of errors are reported. Semi-global alignment can either be between a short pattern and a much longer text (*m* ≪ *n*, e.g. searching a read in a reference genome), or between two sequences of similar length (*m* ≈ *n*, e.g. refining a mapped read). This approach was first introduced by Sellers [1979, 1980], and later termed *semi-global* ^1^ by Gotoh [1999], who also termed *global* alignment. It is implemented in tools such as Parasail [Daily, 2016], SeqAn [Reinert et al., 2017], Edlib [Šošić and Šikić, 2017], and more recently Ish [Stadick, 2025]. When *m* ≈ *n*, semi-global alignment can benefit from adaptive banding methods as developed for global alignment, but this is not the case when *m* ≪ *n*. There, some methods (`parasail, Seqan, Ish`) simply compute the entire *O*(*mn*) DP matrix, while others (`Edlib, Sassy`) often only compute the top *O*(*k*) rows [Myers, 1999]. Thus, these two regimes lead to completely different algorithms. 2248

#### Cost models

A *cost model* defines what constitutes an *error* and the cost associated to each error. This concept originated in the early 1900s with systems designed to detect misspelled names by sound [Odell and Russell, 1918]. In the 1950s, Hamming distance was introduced for binary codes, measuring the number of differing bit positions [Hamming, 1950], or in the context of DNA, the number of mismatches between two equal-length strings. Around a decade later, the Levenshtein distance was formalized [Damerau, 1964, Levenshtein et al., 1966], that allows insertions and deletions alongside substitutions. Importantly, Levenshtein distance uses a *unit cost* model, assigning a cost of 1 to each edit, making it computationally efficient. However, this assumption is unrealistic for large insertions or deletions, as deleting or inserting long segments often represents a single biological event. This led to the development of gap-affine models, where gap opening and extension have different costs [Gotoh, 1982, Altschul and Erickson, 1986, Marco-Sola et al., 2021]. For `Sassy`, we use unit-cost edit distance for its computational efficiency and the assumption that long indels are rare when aligning relatively short patterns.

#### Approximate string matching

As mentioned before, the goal of ASM is to find all matches of a pattern *P* in a text *T* with ≤ *k* errors [Galil and Giancarlo, 1988, Navarro, 2001]. The key distinction from semi-global alignment is that not just the single best match should be reported, but that *all* matches with ≤ *k* errors should be reported. We first discuss *streaming* algorithms, where the text is not known in advance, as opposed to algorithms that pre-process the text *T*. Moreover, we focus on the *k-difference* variant that uses edit distance rather than the *k-mismatch* variant that uses Hamming distance [Chhabra et al., 2025, Fiori et al., 2021, Gottlieb and Reinert, 2025b].

Searching exact matches of patterns became popular through algorithms such as Knuth-Morris-Pratt [Knuth et al., 1977] and Boyer-Moore [Boyer and Moore, 1977]. With the development of different cost models in the 1980s, algorithms were created to detect inexact matches with *k* errors. Initially, *approximate string matching* described comparison of two strings [Ukkonen, 1983, Hall and Dowling, 1980], but later also described searching for a pattern as a substring of a text with ≤ *k* errors [Landau and Vishkin, 1985]. Sellers proposed an *O*(*mn*) time algorithm [Sellers, 1980], which was improved to *O*(*m*^2^ + *k*^2^*n*) [Landau and Vishkin, 1985], and then *O*(*kn*) [Ukkonen, 1985a]. The introduction of bit-parallelism [Baeza-Yates and Gonnet, 1992] led to complexities involving the word size *w*, such as *O*(*k*⌈*m/w*⌉*n*) [Wu and Manber, 1992] in the famous agrep tool, *O*(*mn* log(*σ*)*/w*) [Wright, 1994], and eventually *O*(⌈*m/w*⌉*n*) in Myers’ algorithm [Myers, 1999]. Later optimizations targeted specific scenarios: short patterns or small *k* [Baeza-Yates and Navarro, 1999, Navarro and Baeza-Yates, 1999, Bille, 2011], fixed pattern lengths [Iliopoulos et al., 2001, Ho *et al*., 2017], and periodic texts [Cole and Hariharan, 2002]. Some were optimized for multi-pattern search [Muth and Manber, 1996, Baeza-Yates and Navarro, 1997], though here we focus on a single pattern.

Additionally, many algorithms leverage text pre-processing and indexing, such as text compression [Mäkinen et al., 2003, Kärkkäinen et al., 2000], suffix arrays [Landau and Vishkin, 1986, Manber and Myers, 1993, Huynh et al., 2004], and suffix trees [Ukkonen, 1993]. Others use pre-filtering with n-grams [Owolabi and McGregor, 1988, Jokinen and Ukkonen, 1991, Ukkonen, 1992, Sutinen and Tarhio, 1995, Bingmann et al., 2019], inexact hashing [Yao et al., 2010, McCauley, 2021], heuristics [Koehn and Senellart, 2010, Salmela et al., 2009], or search schemes on top of a birectional FM-index or move-index [Renders et al., 2022, 2024, Gottlieb and Reinert, 2025a, Depuydt et al., 2024, Renders et al., 2025]. Such methods thus implement completely different algorithms than the streaming-based method that we focus on in this paper, and are suitable for different applications.

#### SIMD parallelism

*Overall, the complexity of index-free methods did not improve beyond Myers’ O*(⌈*m/w*⌉*n*). Practical speedups emerged with larger SIMD word sizes with *W* = 256 or *W* = 512 bits. Improvements then involved optimal utilization of *W*. For example, BGSA [Zhang et al., 2019] uses inter-sequence parallelism to compare multiple sequences to the same pattern, since intra-sequence parallelism is limited when *m* ≤ *W* [Zhang et al., 2019]. An alternative approach is taken by A*PA2 [Groot Koerkamp, 2024], where the dependency between SIMD lanes is broken by tiling them diagonally. Yet another approach is taken by SeqMatcher [Espinosa et al., 2024], where AVX512 instructions are used to effectively use 512-bit integers. In contrast, Sassy splits the text into 4 chunks that are processed in parallel, somewhat similar to Farrar’s striped method [Farrar, 2006] and as also used by SimdMinimizers [Groot Koerkamp and Martayan, 2025], for a complexity of *O*(*m*⌈*n/W* ⌉). This way, intra-sequence parallelism is maximized.

#### Mapping

In modern applications, the text is often an assembled genome of many gigabases, and the number of patterns (reads) to be searched is very large. This means that index-free methods are infeasible, and in practice, *mappers* drop the guarantee to find all matches in favour of speed. Thus, we consider mapping to be approximate^2^ ASM.

In the 1980s, with increasing sequence availability and the release of GenBank [Bilofsky et al., 1986] previous exact methods were no longer fast enough. Early mapping methods performed *exact* substring matches between the pattern *P* and database sequences, beginning with Wilbur and Lipman [1983] and followed by others using similar approaches [Lipman and Pearson, 1985, Ning et al., 2001, Wu and Watanabe, 2005] such as BLAST [Altschul et al., 1990].

Some methods controlled sensitivity based on the pigeonhole principle [Weese et al., 2012], while others tried to identify similar regions between *P* and the database sequences through spaced seeds [Ma et al., 2002, Rumble et al., 2009] or locality-sensitive hashing [Buhler, 2001]. As sequences got longer, also the number of seeds increased, leading to algorithms that reduced the number of seeds being stored, such as minimizers (e.g. Minimap2) [Li, 2018, Jain et al., 2020], strobemers (e.g. StrobeAlign) [Tolstoganov et al., 2024], or by hashing subsequences instead of substrings (e.g. SubseqHash2) [Li et al., 2024].

However, in benchmarks these mappers do not detect all mapping locations: they can achieve over 99% sensitivity but not full coverage [Banović Ðeri et al., 2024], and their performance heavily depends on parameter settings such as the seed length [Oliva et al., 2021].

#### Applications of ASM

The earliest methods for ASM, developed in the 1980s, proved directly useful for biological problems. For example, Myers [1986] used ASM to find a 16-nucleotide binding site of the LexA protein in a 48 kb virus genome. Today, ASM supports diverse applications, including read demultiplexing [Cheng et al., 2024], genome polishing [Tonkin-Hill et al., 2020], and CRISPR off-target searching [Chaudhari et al., 2020].

We focus on the latter due to its clinical relevance. CRISPR and its associated Cas proteins form an adaptive immune system in bacteria and archaea, evolved to defend against foreign nucleic acids such as bacteriophage and plasmid DNA [Mojica et al., 2009]. In this system, foreign DNA is precisely cut using a template called single guide RNA (sgRNA). When the target DNA is flanked by a protospacer adjacent motif (PAM) — for example, 5’-NGG-3’ in *Streptococcus pyogenes* — the CRISPR-Cas complex binds and cleaves the DNA, thereby neutralizing the invader. By modifying the sgRNA sequence, the CRISPR-Cas system can be programmed to cut virtually any DNA sequence. This technology has been applied to treat genetic diseases [Ledford, 2019], enhance crop traits [Jaganathan et al., 2018], and engineer microorganisms [Shapiro et al., 2018]. For an in-depth review, see Koonin and Makarova [2019]. Notably, on May 15th, 2025, CRISPR was used for the first time as a personalized treatment for a baby with carbamoyl phosphate synthetase 1 (CPS1) deficiency, a rare and life-threatening condition [Musunuru et al., 2025].

When CRISPR is engineered to target a specific sequence it is crucial that no other, unintended sequences are cut. This is called *off-target* cutting. Hence, computational tools to screen for such off-target sites have been developped. These include Cas-OFFinder [Bae et al., 2014], CRISPRitz [Cancellieri et al., 2020], SWOffinder [Yaish et al., 2024], and CHOPOFF [Labun et al., 2025], with the latter two representing the current state-of-the-art.

While CHOPOFF is much faster than SWOffinder, it requires a time-consuming step of building an index before searching. Given that human genetic variation affects off-target profiles [Scott and Zhang, 2017], we argue that with the advancement of personalized CRISPR therapies, there is a need for fast, index-free tools that are user friendly and robust to ambiguous bases.

## 2. Methods

We now describe our tool, Sassy. We start with some brief notation. Throughout the paper, we assume that we are given a *pattern P* of length *m* := |*P* |, and a *text T* = *t*_0_ … *t*_*n−*1_ of length *n* := |*T* |, which are both strings over an alphabet Σ of size *σ*:= |Σ|. We write *T* [*i* … *j*] := *t*_*i*_ … *t*_*j−*1_ for a right-exclusive substring of *T*, and we use *d*(*P, T* [*i* … *j*]) for the edit distance between *P* and *T* [*i* … *j*]. We write rev(*T*) := *t*_*n−*1_ … *t*_1_*t*_0_ for the reverse of *T*, and for DNA sequences, we define the *complement* comp(*T*) as the sequence where each base is replaced by its complement (A ↔ T and C ↔ G, extended to IUPAC as well). The *reverse complement* is then rc(*T*) := rev(comp(*T*)).

### 2.1 Approximate String Matching

We define approximate string matching following Navarro [2001], but restrict ourselves to edit distance only.

#### Definition 1

(Approximate String Matching, ASM.) Let *P* be a *pattern* of length *m* := |*P* |, and let *T* be a *text* of length *n* := |*T* |. Further, let *k* ∈ 𝒩_*≥*0_ be the maximum number of errors allowed. The problem of *approximate string matching*, search(*P, T, k*), is to find *all* end positions *j* ∈ {0, …, *n*} in the text such that there exists an *i* ∈ {0, …, *j*} such that the edit distance *d*(*P, T* [*i* … *j*]) between *P* and *T* [*i* … *j*] is at most *k*.

#### What is a match?

As defined above, a *match* is a position *j* in the text where an alignment of cost ≤ *k* ends. In practice, one might rather care about all *substrings* of *T* that have edit distance ≤ *k* to the pattern, i.e., all tuples (*i, j*) such that *d*(*P, T* [*i* … *j*]) ≤ *k* [Sellers, 1980, *Definition 1]. Or, even more exhaustively, one could consider all alignments* of *P* to substrings of *T*, where an alignment is a specific sequence of edits transforming *P* into *T* [*i* … *j*].

In `Sassy`, we choose the first option: we find all end positions, and then do a *single* traceback for each of them.

#### When do we have a match?

In practice, one is usually not quite interested in *all* matches. In particular, if there is an exact match ending in position *j*, all positions in *j* − *k* to *j* + *k* will have a cost ≤ *k* (see Figure 2). Thus, there are numerous options for which matches to report:

1. **All** Report (matches ending in) *all* end positions with cost ≤ *k*.
2. **Single best** Report only a *single best* end position (if ≤ *k*). Supported by `Seqan`.
3. **All best** Report *all* positions where a match of *globally optimal* cost ends (if ≤ *k*) as done in semi-global alignment as defined by Sellers [1980] and supported by `Edlib` and `Seqan`.
4. **Non-overlapping** Report only end positions that are at least (roughly) *m* apart. In Sassy, we take a different approach, that we argue is more principled:
5. **Local minima** Report only *rightmost local minima* ≤ *k*.

This is similar to the idea of Sellers [1980] to report all substrings *T* [*i* … *j*] that can not be shrunk nor grown into a substring with lower edit distance, with the difference that we only report end positions, and that we only report a single match for each plateau of local minima.

#### ASM is not reverse-invariant

We note here that when reporting end positions, it is typically hard to guarantee that the matches reported by search(*P, T, k*) are in one-to-one correspondence with those reported by search(rev(*P*), rev(*T*), *k*), since the number of (local/global minima) end positions ≤ *k* can differ in the forward and reverse case, as exemplified in Figure 2. Reporting *all* matching substrings *T* [*i* … *j*] would avoid this, but neither `Sassy` nor other tools do this in practice.

When searching reverse-complements is enabled, `Sassy` *is* invariant to reverse complements of the text: we search *P* in *T* and rc(*T*)^3^, so that both searches are in the natural direction of the pattern and searching *P* in rc(*T*) and rc(rc(*T*)) = *T* gives the same result. Searching rc(*P*) in *T* or rc(*T*) *would* change the set of matches.

#### Traceback

Given the set of end positions that define a match, we can run a traceback from each of them to obtain both the position in the text where the match starts, and the corresponding alignment. `Sassy` simply recomputes the part of the DP matrix preceding each end position and traces back through that. The traceback greedily chooses matches and substitutions if possible, and then falls back in deletions and insertions, in that order.

### 2.2. Bitpacking and SIMD tiling

Figure 1 shows how `Sassy` applies both Myers’ bitpacking [Myers, 1999] and SIMD. Using bitpacking, a *block* of *w* = 64 states of the DP matrix can be computed in parallel. Whereas other methods typically tile these bitvectors in the direction of the pattern, we tile them in the direction of the text. Pseudocode of the main search function of Sassy can be found in Appendix A.

**Fig. 1.**
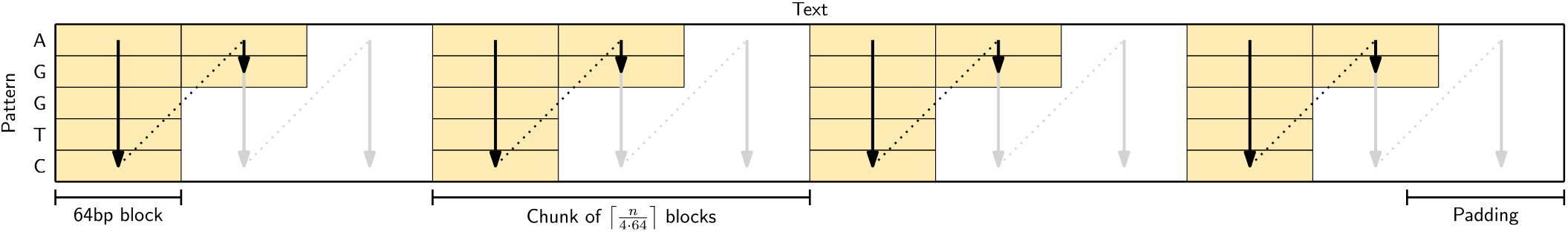
The tiling strategy used by Sassy. The text is first split into word-size blocks of 64 bases. Then, the list of blocks is split into 4 chunks that are processed in parallel, with one SIMD lane per chunk. The text is implicitly padded as needed. Within a chunk, filling the matrix proceeds block-by-block. For each block, all (up to, see Section 2.3 and Figure 3) *m* = |*P* | rows are computed before proceeding to the next block. Each chunk is extended into the succeeding chunk as long as there is a sufficiently good “in progress” alignment (not shown).

**Fig. 2.**
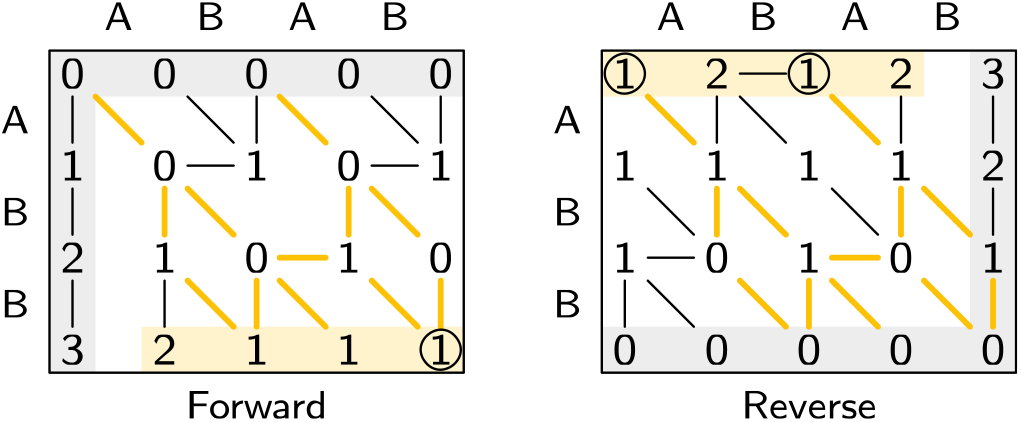
Approximate string matching. An example of finding all occurrences of ABB in ABAB. On the left, the forward search initializes the top and left of the matrix (shaded in grey). Then, it shows all optimal paths to each state. On the bottom, the final distances are highlighted, and all optimal alignments of cost 1 are highlighted in yellow. By default, Sassy only starts a trace in the circled 1, a rightmost local minimum. The right figure shows the reverse alignment, where the matrix is filled from the bottom right to the top left. Note that the set of optimal alignments is the same, but that the number of local minima (1 vs 2) and global minima (3 vs 2) both differ.

#### SIMD

We use 256 bit SIMD widths using either AVX2 or two parallel NEON registers to compute 4 *lanes* of 64 bit words in parallel. We avoid dependencies between SIMD lanes by splitting the text into 4 chunks of ⌈*n/*256⌉ blocks each. Each lane then processes one chunk: we iterate over the 64 character blocks of each chunk, and for each block compute the *m* rows of the matrix. As with Farrar’s striped method [Farrar, 2006], there may be some “patching up” to do when a good alignment crosses the boundary between two chunks. In our case, we extend each chunk to the right (overlapping with the next chunk) as long as there is a partial alignment of cost ≤ *k* that started inside the original chunk. Especially when the text is long (so that ⌈*n/W* ⌉ ≈ *n/W*), this intra-sequence parallellism is near-optimal.

### 2.3 Early break

The complexity of computing the entire DP matrix as described so far is *O*(*m*(⌈*n/W* ⌉ + ⌈*m/w*⌉)), where the final +⌈*m/w*⌉ accounts for overlaps between chunks. In ASM, we only care about matches with a cost at most *k*, and thus, parts of the DP matrix where values are *> k* can be skipped [Ukkonen, 1983, 1985b, Myers, 1986], as shown in Figure 3. In particular, the edit distance between two uniform random and equal-length DNA sequences is typically around 45% of their length, so that most of the time, the cost of aligning a prefix of length 2 · *k* of the pattern already incurs a cost *> k*. More formally, Chang and Lampe [1992] proved that the number of states with cost ≤ *k* is *O*(*kn*) when searching a random text.

**Fig. 3.**
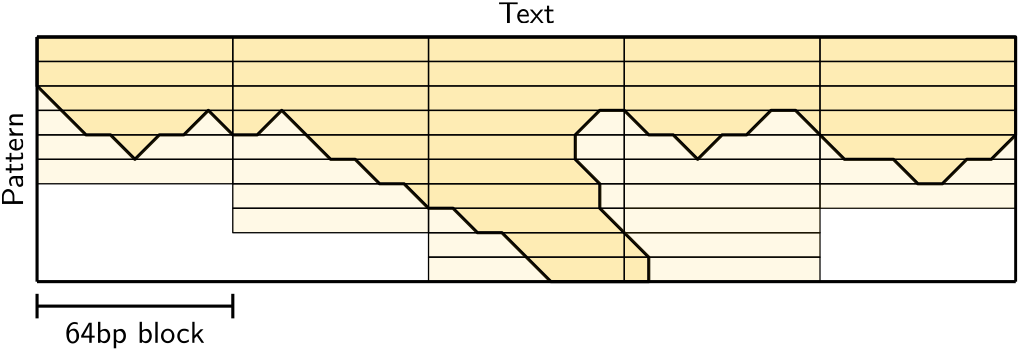
Early break. We are only interested in entries of the DP matrix with value ≤ *k*, as shown in the bold-outlined area. As soon as all entries in a row are *> k*, we can stop processing that block of text, as in the first and second block. Then, we only reach the bottom of the matrix when matches are present, as in the third block. One exception is shown in the fourth block: when there are states at the end of the previous block at distance ≤ *k*, we must continue at least one row beyond that point. Since we use SIMD to process 4 chunks in parallel (not shown here), in practice we continue until the values in all 4 lanes are *> k*.

Thus, as soon as all *W* columns corresponding to a SIMD vector contain a value *> k*, and additionally there are no remaining states with cost ≤ *k* at the end of the preceding blocks, we can stop with the current four blocks and move on to the next.

### 2.4 Library and Command Line Tool

The main entrypoint of the `Sassy` Rust library is a function `search(pattern, text, k)` that returns one match for each rightmost local minium endpoint. It can optionally return matches for *all* endpoints, and supports reverse complements. The input can be either (case insensitive) ASCII text, simple ACGT DNA, or more general IUPAC-encoded DNA where sequences (*both* the pattern and the text) may contain bases such as N (matching ACTG), Y (matching CT), and R (matching AG). This is handled by selecting a *profile* (Appendix C.1). In case of simple DNA, we provide a function to validate that no non-ACGT bases are present. We provide both C and Python bindings.

We also provide a simple command line tool for searching a sequence in all records of a Fasta file, that can be installed via `cargo install sassy` or `conda install -c bioconda sassy`. Examples of commands are:

~~~
sassy search --alphabet dna --no-rc -k 0 --pattern CAT data.fa
sassy search --alphabet iupac -k 1 -f patterns.fa genome.fa.gz
sassy crispr -k 5 --guide guides.fa ref.fa
~~~

The first searches for an exact match of `CAT` in all records of `data.fa`. The second searches each record of `patterns.fa` in `genome.fa.gz`, while allowing up to 1 error and also searching the reverse-complement text. The last searches each of the guides in `guides.fa` against `ref.fa` while allowing at most 5 errors. Here, PAM sequences must match exactly and the preceding sgRNA can contain up to 5 errors.

Whereas the library is single-threaded, the command line tool maintains a queue of (*P, T*) tuples that are distributed (in batches) over all threads. Further details on the implementation can be found in Appendix C.

## 3. Results

We compare `Sassy` against Edlib [Šošić and Šikić, 2017] in two metrics: the throughput of searching random DNA sequences without matches (the number of text bases processed per second), and the throughput of finding and tracing matches (the number of matches that can be found and processed per second). Section 4 shows specific applications of `Sassy`. The code and data for the benchmarks can be found in the `evals` directory at https://github.com/RagnarGrootKoerkamp/sassy. These experiments were run on an Intel Core i7-10750H with 6 cores, 12 threads, AVX2 support, and running at a fixed frequency of 3.6 GHz.

### Throughput of text searching

In Figure 4 and Figure 5, we compare the text throughput of searching with `Sassy` and `Edlib`. Each data point is the average of searching 1000 random DNA pattern of length *m* in a random text of length *n*. Figure 4 compares searching patterns of varying length (20 ≤ *m* ≤ 1000) in a long text (*n* = 10^5^) for varying error thresholds (0 ≤ *k* ≤ 50 = 0.05 · 1000), while in Figure 5, a pattern of length *m* = 100 is searched in texts of length 150 ≤ *n* ≤ 128 000 for *k* ∈ {3, 20}.

**Fig. 4.**
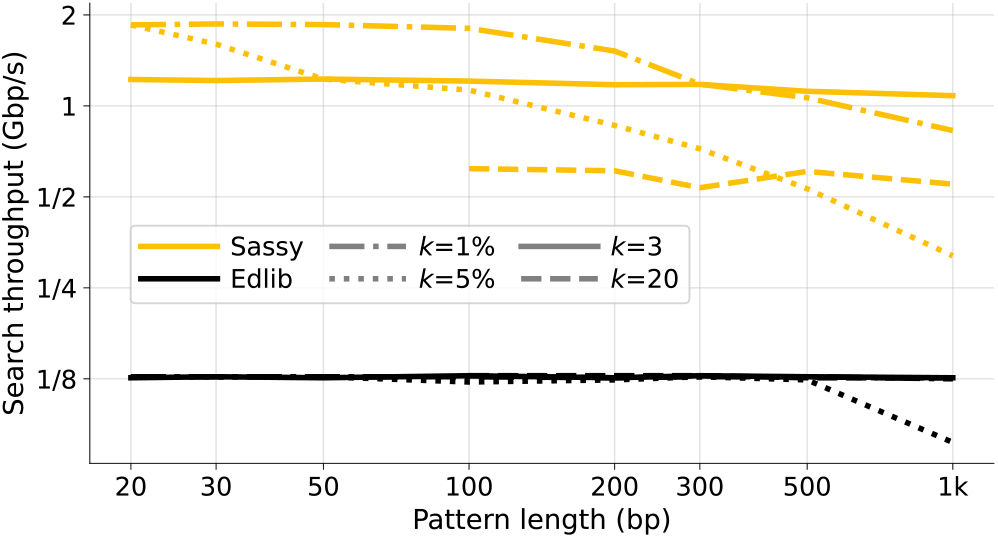
Throughput of searching patterns of varying length. The pattern length *m* (x-axis) ranges from 20 to 1000, and the error threshold *k* (line style) is either fixed at 3 or 20, or computed as ⌈*m/*100⌉ or ⌈*m/*20⌉. Only points with *m >* 3*k* are shown to avoid spurious matches. All points are computed by averaging over 1000 random patterns and texts of length *n* = 10^5^, and then converting to throughput. Note that this does not include searching the reverse-complement strand. Sassy has up to 10× higher throughput than Edlib when *k* is small.

**Fig. 5.**
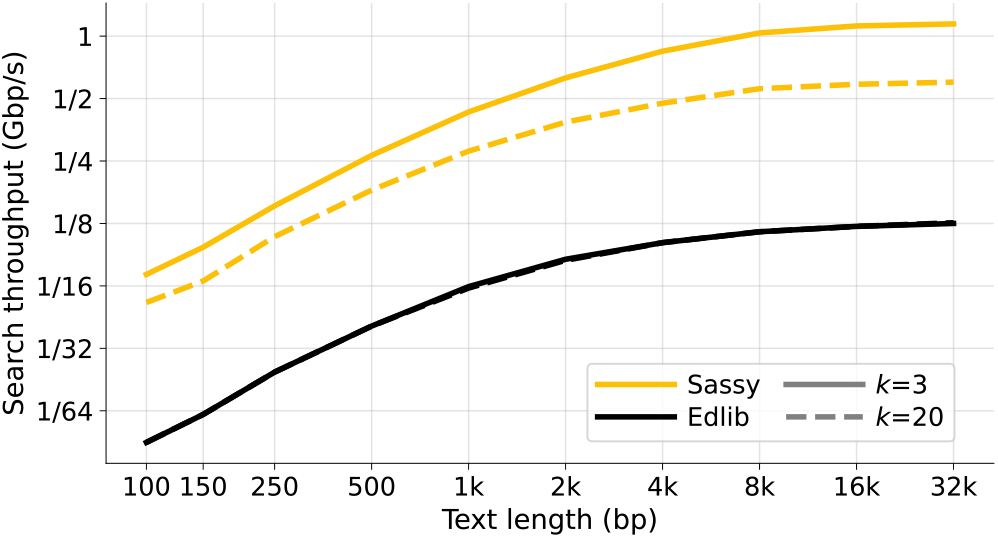
Throughput of searching texts of varying length. We search a pattern of length *m* = 100 against texts with length varying from *n* = 150 to *n* = 128 000 bp, with *k* ∈ {3, 20}. All points are computed by averaging over 1000 random texts and then converting to throughput. Note that this does not include searching the reverse-complement strand. While Sassy is consistently faster than Edlib, its relative advantage is smaller for shorter texts.

We exclude `Seqan`, since it is consistently outperformed by `Edlib`. A comparison against `parasail`, an affine-cost aligner, can be found in Appendix E. It is around 10× slower than `Edlib` and 100× slower than `Sassy`. `Ish` is an up-to 35% faster re-implementation of parasail, but does not provide Rust bindings. Lastly, Other tools such as `agrep` do not accept FASTA input.

`Sassy` is faster across all *m, n*, and *k*. For short patterns (*m* ≤ 50 bp), `Sassy` has throughput over 1.2 Gbp/s whereas the throughput of Edlib does not exceed 130 Mbp/s. Since both the pattern and text are random, no matches occur, and the early break causes `Sassy` to have complexity *O*(*k*⌈*n/W* ⌉). Indeed, for constant *k* the throughput is independent of *m*, while it decreases when *k* = 0.05·*m*. `Edlib`, on the other hand, has throughput nearly independent of *k*: in nearly all cases we have *k* ≤ 20, so that crossing the first *w* = 64 rows of the DP matrix already incurs a cost *> k*. This matches the complexity of *O*(⌈*k/w*⌉*n*). In Figure 5, we see that both `Edlib` and `Sassy` are faster when searching longer texts and have large constant overhead when searching small texts, in case of `Sassy` due to relatively large overheads between text chunks. At text length *n* = 150, `Sassy` is 4× to 6× faster than `Edlib`, which increases to 5× to 9× speedup for longer texts.

The throughput when searching ASCII (slightly faster) or IUPAC (slightly slower) text is within 5% compared to DNA.

### Affine-cost aligners

`Sassy` is over 10× faster than tools implementing affine-cost alignments. For example, `Sassy` needs 4.5 seconds to search a pattern of length 23 in a human genome (with up to *k* = 4 errors, excluding searching the reverse-complement). For the same task, `parasail` [Daily, 2016] in semi-global mode with default cost parameters takes 53 to 69 seconds, depending on the exact configuration (8 or 16 bit values, and SSE4.1 vs AVX2), reporting up to 1.45 GCUPS (`giga` cell updates per second). Ish [Stadick, 2025] with default parameters takes 69 seconds (SSE4.1) to 110 seconds (AVX2). Running `parasail` with the same costs as `Ish` takes 81 seconds to 198 seconds. These methods are slower both because they store larger values (instead of bitpacking), and because they compute two additional *affine layers* of the DP matrix.

We propose that fast edit distance alignments could be used to identify candidate regions which can then be refined with affine-cost methods.

### Throughput of tracing

In Figure 6, we show the throughput of finding matches. This includes the time to locally compute all rows of the matrix (rather than just the top *O*(*k*) rows), the time to recompute the matrix region containing the match, and the time for tracing through the filled matrix. We use the same setup as in previous experiments, and “plant” a single match at the end of the text. We then compared the run time of the same text with and without match.

**Fig. 6.**
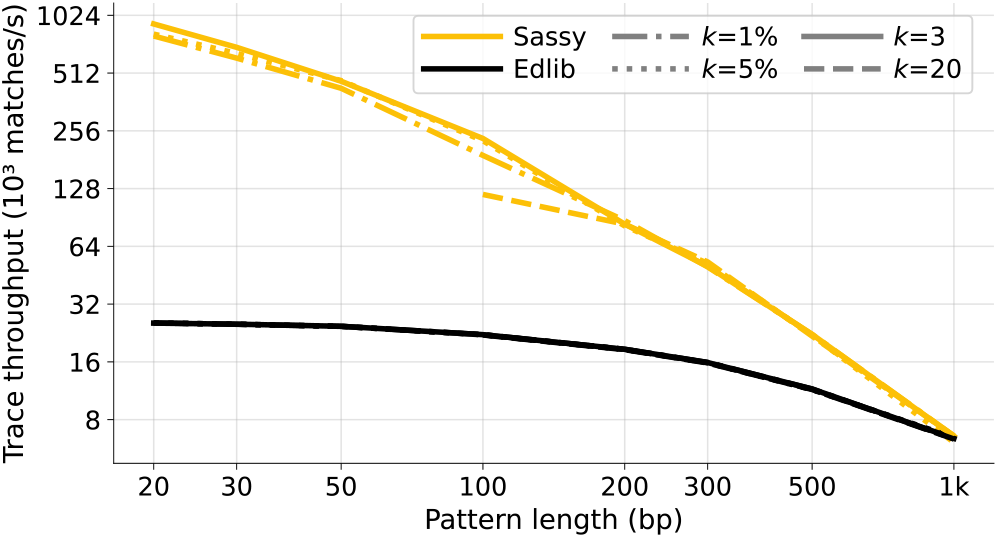
Throughput of finding matches. When the text contains a match, this causes a larger part of the matrix to be computed since the early break does not trigger. Later, this part of the matrix is recomputed in full and stored in *O*(*m*^2^*/w*) words of memory so that a traceback can be done. To measure the total time it takes to process a match, we use the same setup as in fig. 4 with the addition of a single copy of the pattern planted at the end of the text. We measure the time difference with the version without match and report the corresponding throughput.

Placing the match at the end avoids triggering the dynamic reduction of *k* in `Edlib`: Since `Edlib` performs semi-global alignment and only searches matches with the minimum edit distance, it reduces *k* whenever it finds a match with cost *< k*. If the match were placed earlier in the text, this reduction would confound the measurement by timing both the tracing time and its *k* reduction strategy. By placing the match at the end of the text, we largely isolate the tracing cost from the *k* reduction strategy.

For short patterns, `Sassy` is over 10× faster per match than `Edlib`. For longer patterns, `Sassy’s` throughput goes down quadratically as it naively computes the *O*(*m*^2^*/w*) words to fill the part of the matrix where a match is. `Edlib` does not slow down as much, likely due to *O*(*ns*) banding, but is nevertheless still slower than `Sassy` for patterns up to length *m* ≤ 1000.

## 4. Application

### Finding CRISPR off-targets

Finding short sequences has many important applications, with CRISPR off-target searching being of particular recent interest [Musunuru et al., 2025]. SWOffinder [Yaish et al., 2024] and CHOPOFF [Labun et al., 2025] are currently among the fastest and most accurate tools for identifying off-target sequences. We extended `Sassy` with CRISPR off-target searching, enabling the search for PAM motifs—such as the Cas9 NGG motif—preceded by a guide RNA (gRNA) sequence [Ran et al., 2013].

Unlike SWOffinder, `sassy crispr` reports at most one match for every location where the PAM has an exact match.

### 4.1 Searching 61 guide RNAs in the human genome

First, we briefly summarize the algorithms used by the benchmarked tools. SWOffinder uses Smith-Waterman alignment to fill the entire *m*×*n* matrix, identifying all end positions with ≤ *k* edits. It then filters these alignments to only those with at most 1 indel [Yaish et al., 2024]. CHOPOFF takes a different approach by first indexing all PAM locations in the target genome and storing their prefixes i.e. the sgRNA flank. Then they precompute all the edit distance paths, within *k*, through these prefixes. This allows instant lookup of sgRNA sequences with at most *k* edits of previously identified PAM sites. These indexes range from 3.2Gbp (*k* = 0) to 4.5Gbp (*k* = 4) for the human genome. The prefix edit paths are shipped with CHOPOFF for *k* ≤ 4. However, computing this for *k* = 5 did not finish in 10 hours. `Sassy’s` algorithm is similar to its regular search, but with an additional filter prior to traceback: when the user requests the PAM sequence to be unmutated, then the traceback is only performed when the exact PAM is present.

To compare `Sassy` to the other off-target search tools we used the benchmark from the CHOPOFF paper [Labun et al., 2025]. This searches 61 guide sequences with the NGG PAM against the human genome. Experiments were run on a Intel(R) Xeon(R) Gold 6240, using 16 threads for each job. Results are in Table 1.

**Table 1.** Time (mm:ss) to search 61 sgRNAs in Human genome CHM13 using 16 threads, and number of matches. For CHOPOFF the time to build the index is shown in parentheses, and was terminated after 10 hours for *k* = 5. All tools require exact matches of the 3 bp PAM sequence. SWOffinder reports more matches for small *k* because it searches with the PAM sequence at the *front* of the pattern, sometimes resulting in multiple matches for each match of the PAM. For *k* = 5, it has *fewer* matches, because it only allows each match to have up to 1 indel.

| $k$ | Sassy | | CHOPOFF | | Swoffinder | |
| --- | --- | --- | --- | --- | --- | --- |
|  | Time | Matches | Time | Matches | Time | Matches |
| 0 | 0:19 | 62 | 0:18 (19:47) | 62 | 40:21 | 62 |
| 1 | 0:20 | 127 | 0:19 (23:53) | 127 | 42:07 | 155 |
| 2 | 0:23 | 1,284 | 0:19 (23:59) | 1,284 | 39:54 | 1,411 |
| 3 | 0:28 | 32,033 | 0:26 (23:39) | 32,033 | 40:34 | 35,434 |
| 4 | 0:30 | 405,401 | 2:23 (23:56) | 405,401 | 41:38 | 471,395 |
| 5 | 0:44 | 4,093,387 | – | – | 40:43 | 1,476,640 |

For large values of *k*, `Sassy` outperforms both competitors by a wide margin. In fact, for *k* ≥ 4, `Sassy` is more than 100× faster than SWOffinder and over 4× faster than the index-based CHOPOFF. Notably, `Sassy` completes the *k* = 5 search in just 44s, whereas CHOPOFF’s index building for *k* = 5 exceeded 10hours (and was therefore omitted). For smaller values (*k* ≤ 3), `Sassy` and CHOPOFF have comparable performance, with `Sassy` trailing by only a few seconds. We do note that CHOPOFF is faster when there are substantially more sgRNA patterns, as most time is spent on loading the index, which is not parallellized over multiple threads.

We note that full support for IUPAC bases, as `Sassy` and CHOPOFF have, is important for this application, since human genome assemblies may not always be fully resolved—see Appendix D.

Thus, `Sassy` is an extremely fast tool that does not require building an index, making it ideal for personalized, reference-free, off-target screening.

## Discussion

`Sassy` solves approximate string matching, and allows fast searching for short DNA sequences without using an index. The main algorithmic novelty is to use *horizontal* bitpacking of deltas, and intra-sequence parallellism using SIMD (Figure 1), leading to a complexity of *O*(*k* · ⌈*n/W* ⌉) when searching random text. This improved complexity allows searching text at nearly 2 Gbp/s, and up to 15× speedup over `Edlib`.

Practically speaking, `Sassy` is a simple-to-use tool with many applications. Since `Sassy` is index-free, it easily supports IUPAC characters in both the pattern and text. It is significantly faster than other index-free methods for searching CRISPR off-target matches, and is being integrated into other tools such as CRISPRapido^4^, which uses `Sassy` as a pre-filter for off-target detection with a more fine-grained (affine-cost) cost model, and Barbell [Beeloo et al., 2025], a demultiplexer for Nanopore reads.

It is left to the user to choose a suitable edit distance threshold *k* that captures all biologically relevant matches, and to post-process and/or refine the matches with a more accurate affine or position-specific cost model if needed.

### Limitations and future work

When text that is searched is short (*n* ≤ 1000 or so, and in particular for *n* = 150), `Sassy` fails to reach its maximum throughput (Figure 5), because the overhead of initializing each search is relatively large and there is a lot of overlap between adjacent text chunks. This is particularly relevant when searching barcodes in reads, and motivates ongoing work on Sassy2, which searches batches of multiple patterns at once.

`Sassy` is primarily designed to search patterns with length *m* ≤ 100 or so, and includes a quadratic *O*(*m*^2^*/w*) component in both the initial filling of the matrix (Figure 3) and the traceback, whereas a banded *O*(*mk/w*) approach would be sufficient in theory.

Lastly, `Sassy` only provides limited benefit when a *static* text is searched with many (at least 100 to 1000) patterns: in that case, search schemes for approximate string matching [Renders et al., 2022, 2024, Gottlieb and Reinert, 2025a] that use a bidirectional index, such as Columba [Depuydt et al., 2024, Renders et al., 2025], should be preferred instead.

## Acknowledgments

We thank Erik Garrison for encouraging us to build `Sassy`, and Rob Patro for feedback on this paper. We thank Seth Stadick for help with the evaluation of `parasail` and `Ish`.

RB is financed by ZonMw [541003001]. RGK is financed by ETH Research Grant ETH-17 21-1 to Gunnar Rätsch.

## A. Pseudocode

### Alg. 1

**Pseudocode** for Sassy’s main Search function, that takes as input the pattern *P* of length *m* and text *T* of length *n*, and returns a list of end-positions in the ext where alignments of cost ≤ *k* end. Some of the operations operate on SIMD registers such as [0, 0, 0, 0]. ComputeBlock applies Myers’ bitpacking algorithm to SIMD registers. Details of Eq, AllAboveK andfind End positionsAt MostKcan be found in Appendix C.

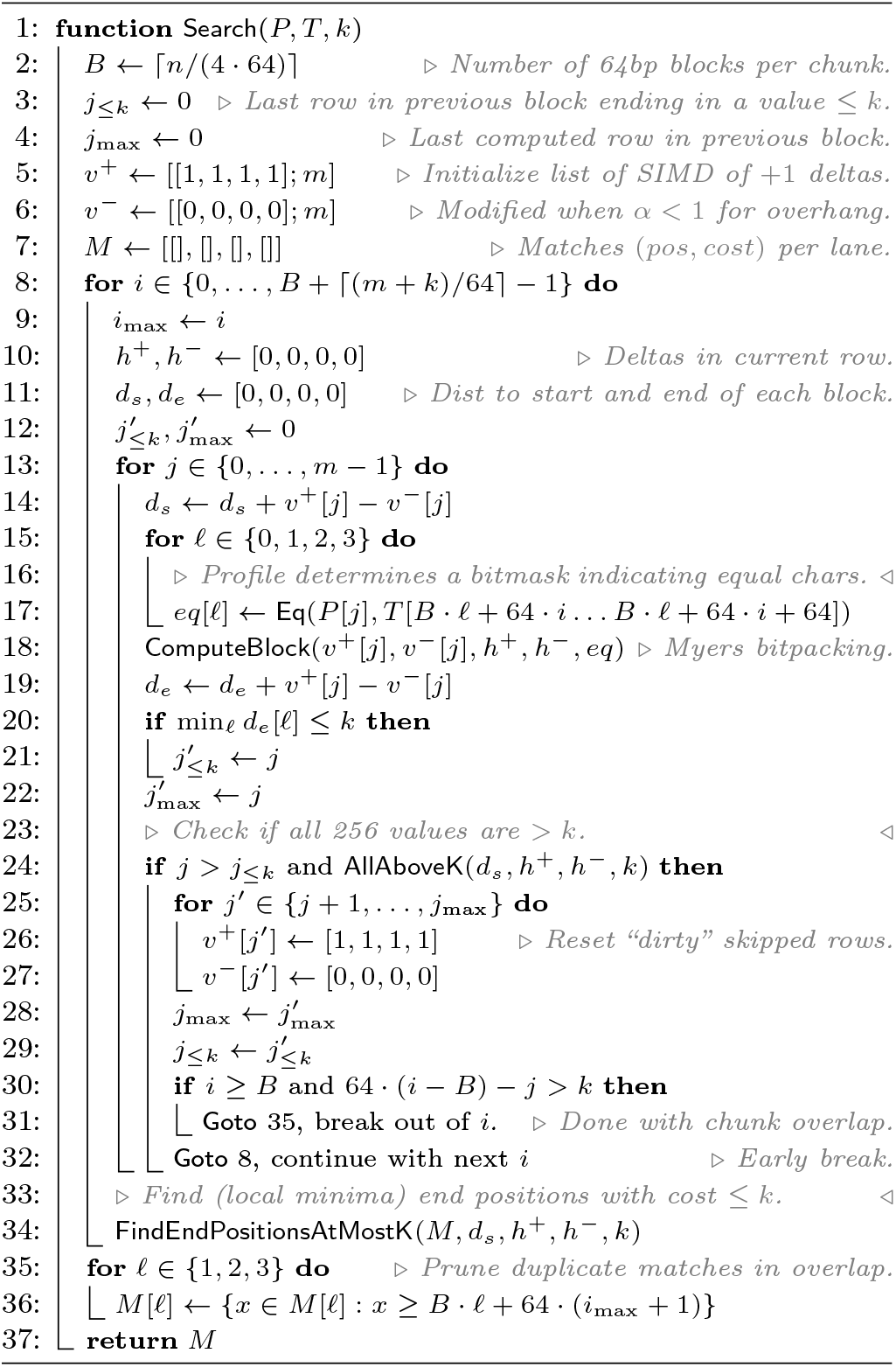

## B. Overhang alignments

In some applications, it is useful to not only find occurrences of the pattern that are fully contained in the text, but also those that extend beyond the text. For example, this is the case for reads that may only partially contain a barcode, or in case of fragmented assemblies [Abramova et al., 2024]. Following Abrahamson [1987], we call these *overhanging* matches.

### Definition 2

(Overhanging match) Given pattern *P* of length *m* and text *T* of length *n*:

1. there is a *left-overhanging* match when a suffix *P* [*l* … *m*] matches a prefix of *T*,
2. there is a *right-overhanging* match when a prefix *P* [0 … *m* − *l*] matches a suffix of *T*.

In either case, each of the *l* overhanging (unmatched) characters of *P* incurs a cost of 0 ≤ *α* ≤ 1, for a total *overhang cost* of ⌊*l* · *α*⌋.

Figure 7 shows an example with *α* = 0.5 (Sassy’s default) with three matches of cost 1: a left-overhanging match, a non-overhanging match, and a right-overhanging match. When *α* = 1, this corresponds to semi-global alignment and ASM, whereas *α* = 0 corresponds to *overlap* alignments.

**Fig. 7.**
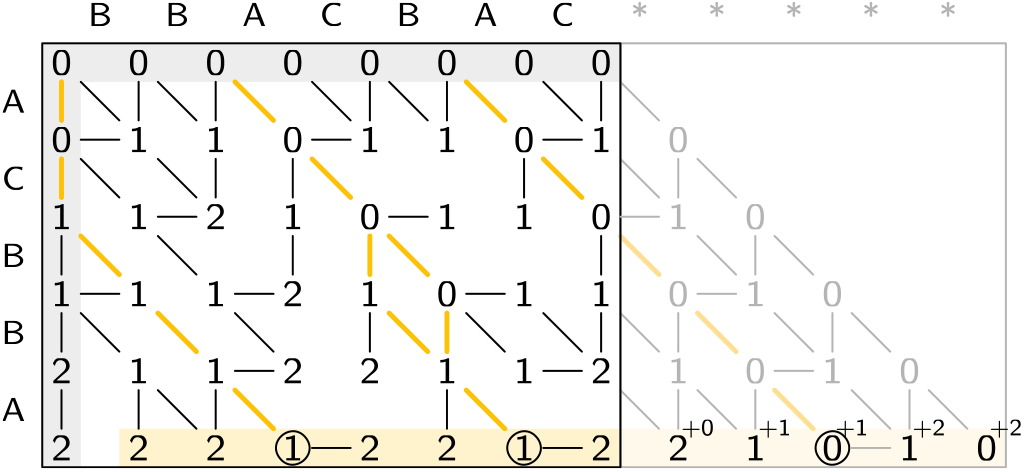
**Overhang alignment** of ACBBA against BBACBAC, with overhang cost *α* = 0.5. On the left, the state in row *j* is initialized with cost ⌊*jα*⌋, and a left-overhanging alignment (highlighted) is found where BBA matches and AC extends beyond the text for a cost of ⌊2*/*2⌋ = 1. On the right, the text is padded with *m* wildcard symbols, so that the costs on the right side of the matrix are replicated in the bottom row. Then, for each extended state in columns *i > n*, ⌊(*i* − *n*)*α*⌋ is added to the cost. This finds a right-overhanging alignment where AC matches and BBA extends beyond the text for a cost of ⌊3*/*2⌋ = 1.

`Sassy` only needs a slight modification to find overlapping matches: the left of the DP matrix is initialized with cost ⌊*iα*⌋ instead of *i*, and on the right we extend the text with *m* “wildcard” symbols, so that the costs in the right column of the matrix are “copied” to the bottom row. Then, we manually add the overhang cost for those extended states. In practice, we use the IUPAC alphabet for this, and simply append N characters.

## C Implementation Notes

We now provide some more details on the functions used in Algorithm 1.

AllAboveK(*d*_*s*_, *h*^+^, *h*^*−*^, *k*). This function takes as input the distance *d*_*s*_ to the start of 4 blocks, a threshold *k*, and the bit-encoded horizontal differences in each block. It then checks if the represented values in all blocks are *> k*. In practice, this function is slightly slow, and we call it not every row (as shown in the pseudocode), but only at most every 8 rows.

It processes each lane *ℓ* independently. First, we *pack* the bits of *h*^+^[*ℓ*] and *h*^*−*^[*ℓ*] into *p* (`using _pext_u64` available in BMI2) to remove positions where the delta is 0 and both have an indicator bit of 0. Then, a 1 in *p* indicates −1 and a 0 indicates +1 (with +1 padding at the end). We split the 64-bit value *p* into bytes, and do a precomputed table lookup that gives the total delta across the byte, and the minimum prefix sum in the byte. The we do a rolling sum over the bytes to compute the minimum overall prefix.

FindEndPositionsAtMostK(*M, d*_*s*_, *h*^+^, *h*^*−*^, *k*). This function again works lane-by-lane. Unlike AllAboveK, this function is only called at most once per block of text, when the iteration over *j* reaches the end of the pattern. Thus, it is less performance critical, and we implement it by simply iterating through the 64 bits of the bitmasks and keeping a rolling sum for the current score in each column. Then, each time a value ≤ *k* is seen, the index of the text and the corresponding cost are pushed to the list of matches.

### C.1. Profiles

The job of a *profile* is to take a single character *P* [*j*] of the pattern and a slice of 64 text characters, and determine a bitmask Eq(*P* [*j*], *T* [*x* … *y*]) indicating which of the text characters equal *P* [*j*] [Rognes and Seeberg, 2000].

We recommend having a look at the code, in the `src/profiles` directory of the git repository. Currently we only support AVX2, and thus, this is subject to change.

#### ASCII

For the ASCII profile, we implement this as follows. For each block of text, we precompute a 256 long array of 64-bit words, so that the mask for each ASCII (In fact, ASCII characters are *<* 128, but this way we can handle any raw data.) character of the pattern can simply be looked up. The array is filled by using 256-bit SIMD instructions to compare each byte up to 256 to both the first 32 characters ([`u8; 32`] is 256 bits) and last 32 characters of the text slice, and merging the two 32-bit values.

For efficiency, we first compute a list of all distinct bytes in the pattern, and then only fill table rows corresponding to those bytes.

#### Case-insensitive ASCII

In this case, we first lowercase all text and pattern characters before doing the equality check. This is done by xor’ing the value of all uppercase bytes by 32.

#### DNA

DNA only has 4 characters, and so we precompute a table of size 4. Each ACTG character is encoded into an integer in {0, 1, 2, 3} by first shifting right 1 bit and then only looking at the bottom 2 bits.

Optionally, it can first be checked that the text only contains valid bases in ACTG. This is done by ensuring that each position case-insensitively equals one of ACTG.

#### IUPAC

Here, we start by building a table that maps each IUPAC character to a 4-bit mask indicating which subset of ACTG it matches. Since only letters are allowed input, it is sufficient to only consider the low 5 bits of each input character leaving 32 possible values. This automatically collapses upper and lower case values. We would now like to use a [`u8; 32`] SIMD register as a lookup table (via shuffle instructions), but unfortunately cross-128-bit lane byte shuffles are not supported on AVX2. We work around this: each byte only contains 4 bits of data, and thus, we can merge them, so that byte *i* in a [u8; 16] contains the 4 bits of both *i* (low half) and *i* + 16 (high half). Then, we can use this as a lookup table on the low 4 bits of each text character, and use the 5th bit to select either the low or high half of the returned byte.

From here, we proceed similar to before: we first build a list of characters occurring in the profile. Then we encode each character to its 4-bit representation, and find the text characters that this “intersects” with.

## D. Support for ambiguous bases

Depending on their quality, human genome assemblies can contain over 10% ambiguous bases, as seen in GRCh38 [Nurk et al., 2022]. In search applications with clinical implications, such as CRISPR off-target analysis, it is crucial to report matches in regions containing ambiguous bases (e.g., N), as these indicate sequence uncertainty and may harbor unintended cut sites. To evaluate tool performance in such scenarios, we searched for the sgRNA GGAAGACACACTGGCAGAAANGG with *k* = 0 against a mock sequence where the sgRNA base at position 14 (C) was replaced by N in one version of a text, and by Y in another. Sassy implements the IUPAC profile for CRISPR off-target searches and returned all matches according to IUPAC base pairing. CHOPOFF requires a user to specificy the max number of ambiguous bases (we used `–ambig-max=23`) and did also return all matches. SWOffinder does not have a command line option but a hardcoded boolean flag (default is `false`) which we set to true and recompiled. It did find the N version but not the Y version. Therefore, both `Sassy` and CHOPOFF have IUPAC support, and SWOffinder only supports N with source code modification. This result underscores the importance of selecting tools that correctly handle ambiguous bases in clinically relevant analyses.

## E Comparison with parasail

We compared `Sassy` to `parasail`, a SIMD-based affine-cost aligner [Daily, 2016], and to `Edlib`. To minimize overhead, we used the Rust bindings at https://github.com/nsbuitrago/parasail-rs to call parasail as a library, consistent with our setup for Sassy and Edlib.

Because affine scoring is not directly comparable to edit distance, we approximated edit distance costs by setting `–gap-open=1, –gap-extension=1, match-score=0`, and `mismatch-score=-1`. Given the small range of scores under this configuration, we used 8-bit output (`solution_width=8`).

`parasail` supports diagonal, striped, and prefix-scan vectorization. We used prefix-scan as it was the fastest for increasing pattern and text lengths.

As shown in Figure 8, when searching a text of *n* = 100 000 bp with varying pattern lengths, `Sassy` achieves approximately 10× higher throughput than `Edlib` and 100× higher throughput than `parasail`. Similar trends hold when varying the text length (Figure 9).

**Fig. 8.**
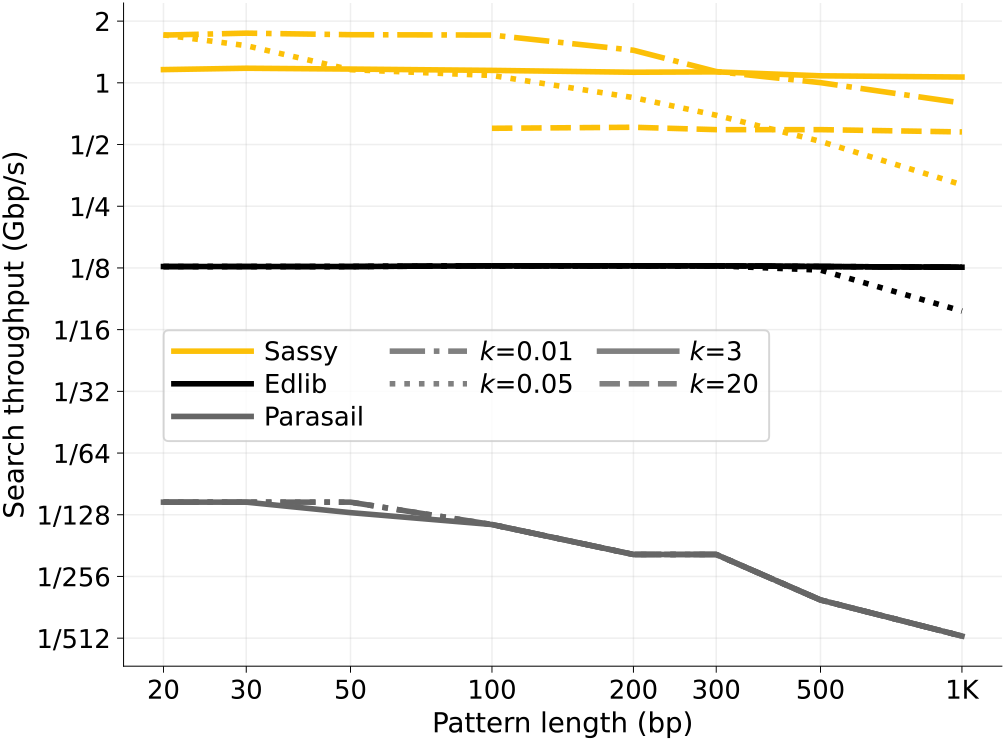
Throughput of searching patterns of varying length. The pattern length *m* (x-axis) ranges from 20 to 1000, and the error threshold *k* (line style) is either fixed at 3 or 20, or computed as ⌈*m/*100⌉ or ⌈*m/*20⌉. Only points with *m >* 3*k* are shown to avoid spurious matches. All points are computed by averaging over 1000 random patterns and texts of length *n* = 10^5^, and then converting to throughput. Note that this does not include searching the reverse-complement strand. Sassy achieves up to 10× higher throughput than Edlib for small *k*, and roughly two to three orders of magnitude higher throughput than parasail. The performance gap with parasail increases with pattern length, exceeding 500× at *m* = 1000.

**Fig. 9.**
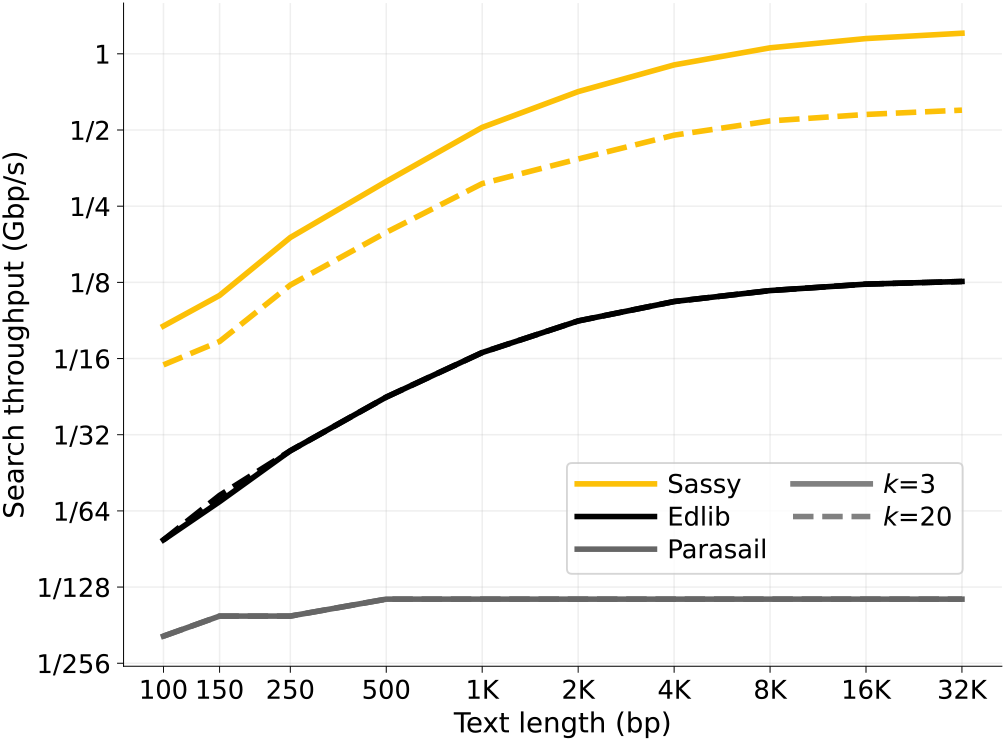
Throughput of searching texts of varying length. We search a pattern of length *m* = 100 against texts with length varying from *n* = 150 to *n* = 128 000 bp, with *k* ∈ {3, 20}. All points are computed by averaging over 1000 random texts and then converting to throughput. Note that this does not include searching the reverse-complement strand. Sassy consistently outperforms Edlib by about one order of magnitude, while parasail remains roughly two to three orders of magnitude slower but less sensitive to text length.

## Footnotes

1 Confusingly, the term semi-global is sometimes also used for different variants of alignment. Parasail [Daily, 2016] uses it for all types of alignment that are not exactly global alignment, while Suzuki and Kasahara [2018] use it for *extension* alignment where the pattern has to match at the start of the text.

2 ASM is *approximate* in the sense that matches are allowed up to *k* errors. Mapping is *approximate* in the sense that it is an approximate algorithm that does not guarantee to find *all* such matches.

3 As an implementation detail, we actually search comp(*P*) against rev(*T*), so that we can avoid taking the complement of *T*. Invariance under complements is trivial.

4 https://github.com/pinellolab/crisprapido

